# A new pipeline for cross-validation fold-aware machine learning prediction of clinical outcomes addresses hidden data-leakage in omics based ‘predictors’

**DOI:** 10.64898/2026.03.12.711429

**Authors:** Marcelo Hurtado, Vera Pancaldi

**Affiliations:** Centre de Recherches en Cancérologie de Toulouse, CRCT, Université de Toulouse, INSERM, CNRS, Toulouse, France; Equipe labellisée Ligue Contre le Cancer

## Abstract

**Motivation:** Machine learning (ML) approaches are increasingly applied to high-dimensional biological data in which features are often dataset-dependent. In many omics workflows, features are computed using information derived from the entire dataset, such as correlations between variables, clustering structures, or enrichment scores. We refer to these as *global dataset features*, defined as features whose computation depends on properties of the full dataset. In such cases, standard validation strategies can fail, especially when evaluating on independent datasets, due to information leakage that leads to overly optimistic performance estimates.

**Results:** To address this challenge, we present *pipeML*, a flexible and modular machine learning framework designed to support leakage-free model training through custom cross-validation (CV) fold construction. *pipeML* enables users to recompute *global dataset features* independently within each CV fold, ensuring strict separation between training and test data, while preserving compatibility with a wide range of ML algorithms for both classification and survival tasks. Using real-world biological datasets, we demonstrate that *pipeML* enables leakage-free model evaluation when *global dataset features* are used. We argue that overestimation of model performance during CV can lead to overoptimistic expectations for validation on independent datasets. By explicitly addressing data leakage and offering a transparent, modular workflow, *pipeML* provides a robust solution for developing and validating ML models in complex biological settings.

**Availability:** The *pipeML* R package as well as a tutorial are available at https://github.com/VeraPancaldiLab/pipeML

**Supplementary information:** Available at *Bioinformatics* online.

## 1 Introduction

Machine learning methods are increasingly applied to high-dimensional biological data to identify predictive signatures and stratify patients. In many biological analyses, however, features are not independent measurements but they are instead derived from relationships across multiple samples. Examples include pathway activity scores, gene set enrichment statistics, transcription factor activity estimates, or cell-state aggregates inferred from gene expression profiles. These features are often computed using correlations, enrichment statistics, clustering structures, or other transformations, learned from the data itself. We refer to such representations as *global dataset features*, defined as features whose computation depends on properties of the entire dataset, including the number of samples, their relationships, or global statistics summaries. The use of *global dataset features* creates important challenges for machine learning model evaluation. When such features are computed on the full dataset prior to model training, information from samples that later appear in test partitions can inadvertently influence the feature representation. As a result, standard cross-validation strategies no longer maintain strict independence between training and validation data, leading to overly optimistic performance estimates. This issue is particularly critical in biomedical applications, where datasets are often small, heterogeneous, and collected across multiple cohorts, making reliable performance estimation essential.

Despite its importance, leakage-aware feature construction is rarely addressed in existing machine learning pipelines, which typically assume that features are fixed and independent across samples. As a consequence, feature construction is frequently treated as a preprocessing step applied before cross-validation, implicitly allowing information from validation samples to influence model training. One possible strategy to avoid information leakage is to learn *global dataset features* using a fully independent dataset and then project them onto the data used for model training and evaluation (Lapuente-Santana, 2021, 2024). While this approach is effective, it requires large, representative, and biologically independent datasets aligned with the target cohort, conditions that are often not met in practice, particularly in omics studies characterized by limited sample sizes, batch effects, and context-specific biological signals.

There is therefore a pressing need for flexible machine learning frameworks that explicitly support fold-aware, leakage-free feature generation while maintaining robust model training, evaluation, and interpretability. To address these issues, we introduce *pipeML*, an R package that provides a flexible and modular framework for leakage-aware machine learning analysis of high-dimensional biological data. *pipeML* supports fold-aware feature construction, enabling *global dataset features* to be recomputed independently within each training fold during cross-validation. This design preserves strict separation between training and validation data, while maintaining compatibility with a wide range of machine learning algorithms for both classification and survival tasks. It integrates feature selection, repeated and stratified cross-validation, model training and validation, benchmarking of several algorithms, hyperparameter tuning, prediction, and interpretation of features through SHAP analyses, all within a unified pipeline. It further supports realistic model evaluation and advanced validation strategies, including cross-dataset generalization such as leave-one-dataset-out analysis (***Figure 1***).

**Figure 1.**
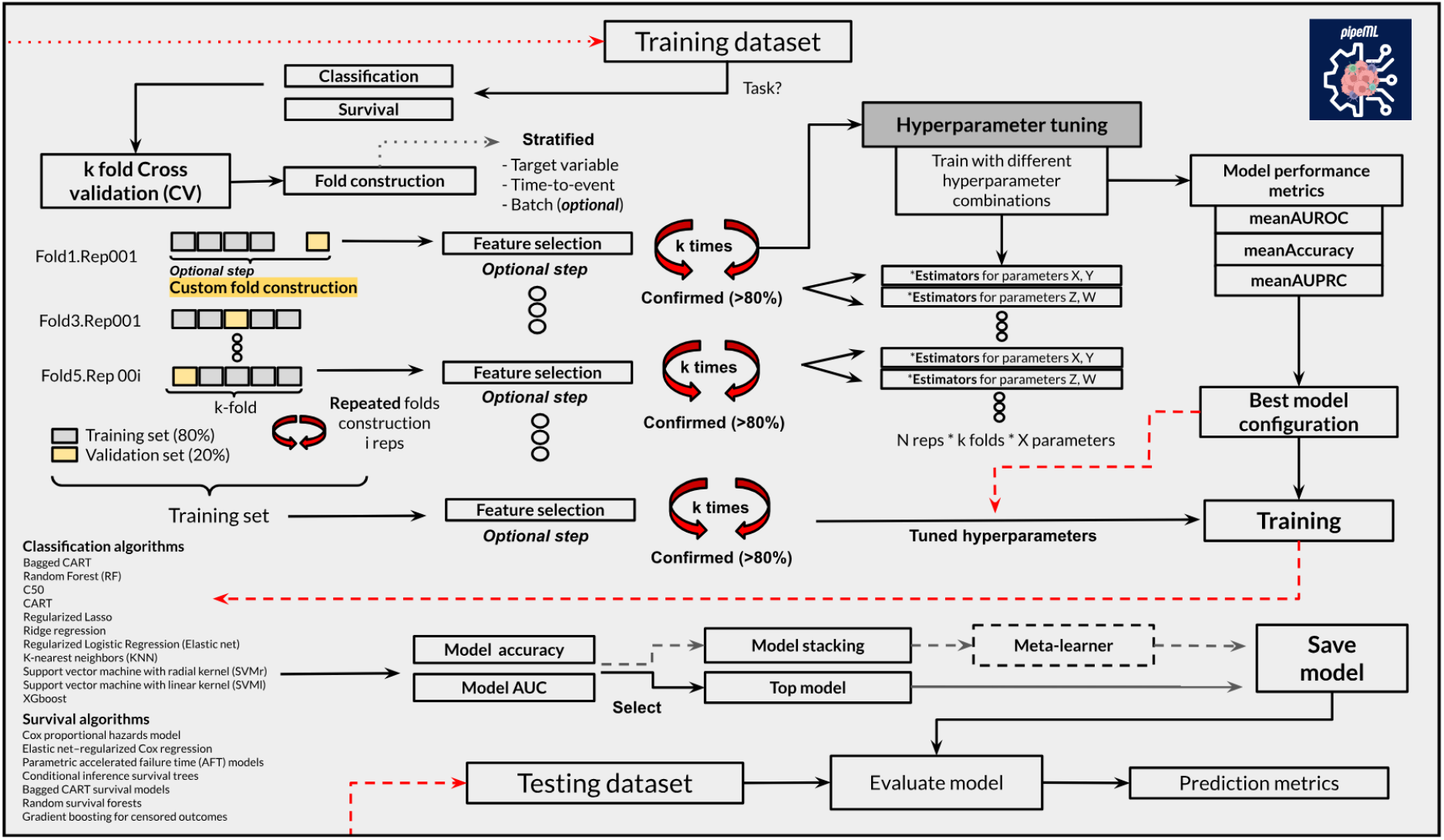
pipeML framework

While several machine learning frameworks, including scikit-learn, AutoML and FLAML provide powerful tools for automated model development, many are primarily developed in Python and do not natively integrate with the R/Bioconductor ecosystem, where a large proportion of bioinformatics tools for omics data analysis are implemented. As a result, researchers working within R-based bioinformatics workflows often face challenges when incorporating leakage-aware machine learning pipelines into their analyses. *pipeML* addresses this gap by providing a framework designed specifically for the R environment, facilitating seamless integration with existing bioinformatics packages.

## 2 Methods

*pipeML* implements a machine learning pipeline for classification and survival tasks, relying on the R packages *caret* (v6.0.94), *tidymodels* (v1.4.1), *parsnip* (v1.3.3) and *censored* (v0.3.3).

### 2.1. Using *pipeML* for binary classification tasks

For classification tasks, we implement a diverse set of classification algorithms that are benchmark on the fly, including tree-based methods, regularized linear models, support vector machines with radial and linear kernels, and instance-based learning.

Model performance is assessed using repeated stratified k-fold cross-validation to ensure robust estimation and preserve class distributions across folds. In this framework, the dataset is partitioned into k folds and the cross-validation procedure is repeated multiple times with different fold assignments. Hyperparameter tuning is performed within the cross-validation loop using grid search. Model selection can be optimized according to a user-defined performance metric, including the area under the receiver operating characteristic curve (AUROC), the area under the precision–recall curve (AUPRC), or classification accuracy, depending on the task.

The current implementation supports multiple classification algorithms, including:

- Bagged classification trees
- Random forest
- C5.0 decision trees
- Regularized logistic regression (elastic net)
- k-nearest neighbors
- Classification and regression trees (CART)
- Lasso regression
- Ridge regression
- Support vector machines with linear and radial kernels
- Extreme gradient boosting (XGBoost)

### 2.2. Using *pipeML* for survival tasks

For time-to-event outcomes, we implement a unified survival modeling framework based on the *parsnip* and *workflows* R ecosystems, enabling consistent training, tuning, and evaluation across multiple survival model families. Models are specified using a censored regression formulation and fitted using appropriate computational engines for each algorithm.

When applicable, model-specific hyperparameters (e.g., regularization strength, mixing parameters, or number of trees) are tuned via grid search within the training data. All models are trained using cross-validation strategies consistent with the main pipeline, ensuring that feature construction and hyperparameter selection are confined to the training folds.

Model performance is then evaluated on held-out data using the Concordance index (C-index), which measures the ability of the model to correctly rank survival times under right-censoring.

The current implementation supports multiple survival algorithms, including:

- Cox proportional hazards model
- Elastic net–regularized Cox regression
- Parametric accelerated failure time (AFT) models
- Conditional inference survival trees
- Bagged CART survival models
- Random survival forests
- Gradient boosting for censored outcomes

### 2.3. Machine learning model selection

The pipeline includes an automated procedure to identify the best-performing model according to a user-defined metric (Accuracy, AUROC, AUPRC or C-index). Models are ranked based on the median performance obtained during cross-validation, and the model with the highest score is selected as the top model. This model can subsequently be used as the final predictive model when model stacking is not applied.

#### Hyperparameter tuning strategy

Hyperparameter tuning is embedded directly within the cross-validation framework to ensure unbiased model selection and prevent information leakage. For each repeated and stratified k-cross-validation fold split, candidate hyperparameter configurations are evaluated exclusively on the training cross-validation folds. When no custom feature cross-validation fold-construction is specified, model-specific tuning grids defined by the caret framework are used, and optimal hyperparameters are selected based on cross-validated performance using the primary optimization metric (Accuracy, AUROC, AUPRC or C-index). When a custom cross-validation fold construction function is provided (e.g., for *global dataset features* generation such as WGCNA as exemplified in this paper), hyperparameter tuning is extended to jointly evaluate both model-level parameters (e.g., number of trees, regularization strength, kernel parameters) and feature-construction parameters. In this setting, each cross-validation fold contains multiple sub-cross-validation folds corresponding to different feature-construction configurations, and all machine learning models are trained and evaluated independently for each configuration. Performance metrics are aggregated across all resamples, and the optimal combination of model hyperparameters and feature-construction parameters is selected based on the median cross-validated metric. Finally, models are retrained on the full training dataset using the selected optimal parameters.

#### Model stacking

*pipeML* optionally implements model stacking to combine predictions from multiple base learners. In this approach, each machine learning algorithm generates probabilistic predictions for the outcome variable. These predictions are then used as input features for a meta-learner, based on logistic regression, which learns how to optimally combine them. It also enables the calculation of feature importance scores based on the contribution of each base learner to the final stacked model.

### 2.4. Model interpretability

Model interpretability is assessed using SHAP (SHapley Additive exPlanations) values (Lundberg & Lee, 2017) to quantify the contribution of individual features to model predictions. SHAP values are estimated using the R package *fastshap* (v0.1.1) (Greenwell B, 2024), which provides a model-agnostic Monte Carlo approximation of Shapley values. For each trained model, SHAP values are computed separately within each cross-validation resample to account for variability in model fitting across training folds. Within each resample, the model is retrained using the optimal hyperparameters identified during model tuning, and SHAP values are estimated for the corresponding hold-out samples.

The *fastshap::explain* function is then applied using the training data of the corresponding fold as the background dataset and the hold-out samples as the evaluation dataset. The SHAP values therefore represent the contribution of each feature to the predicted probability for each individual sample, relative to the model’s baseline prediction. SHAP values obtained across all resamples are subsequently combined to produce a single matrix of per-sample feature contributions. Global feature importance is summarized by computing the mean absolute SHAP value across samples, providing a ranking of features according to their overall influence on model predictions.

For visualization and interpretation, users can then employ the R package *shapviz* (v0.10.3), which enables graphical summaries such as feature importance plots, SHAP beeswarm plots, and dependence plots. These visualizations combine SHAP values with the corresponding feature values, allowing assessment of both the magnitude and direction of feature effects on model predictions.

#### Feature selection

Feature selection can be performed using the *Boruta* R package (v8.0.0). To improve the stability of feature selection, *pipeML* executes it multiple times using different random seeds. In each iteration, Boruta compares the importance of original variables with that of randomly permuted “shadow” features, to determine whether a variable carries information beyond random noise. Features are assigned one of three labels: Confirmed, Tentative, or Rejected. To obtain robust feature importance estimates, results from all iterations are aggregated by computing the median importance of each feature and summarizing the frequency of Boruta decisions across runs. Features are retained if they are classified as Confirmed (or optionally Tentative) in more than a user-defined proportion of iterations.

### 2.5. Performance metrics and threshold-based evaluation

Model performance is evaluated using several threshold-dependent and threshold-independent metrics, derived from predicted probabilities for the positive class. Predictions are first sorted in decreasing order of the predicted probability, and sensitivity–specificity values were computed across all possible classification thresholds. For each threshold, cumulative counts of true positives (TP) and false positives (FP) are obtained using cumulative sums. Sensitivity (true positive rate) is defined as the proportion of correctly identified positive samples relative to all positive samples, while specificity is calculated as one minus the false positive rate (FPR).

From the confusion matrix components (TP, TN, FP, FN), additional classification metrics are computed. Accuracy is defined as the proportion of correctly classified samples among all observations. Precision is calculated as the proportion of true positive predictions among all predicted positives, whereas recall (sensitivity) measures the proportion of true positives among all actual positive samples. The F1 score is computed as the harmonic mean of precision and recall, providing a balanced measure when both false positives and false negatives are relevant. The Matthews correlation coefficient (MCC) is also calculated as a balanced metric incorporating all four confusion matrix components, ranging from −1 (complete disagreement) to 1 (perfect prediction), with 0 indicating random performance.

Threshold-independent performance is assessed using receiver operating characteristic (ROC) and precision–recall (PR) curves. ROC curves are generated from sensitivity and false positive rate values across thresholds, and the area under the ROC curve (AUROC) is calculated using the trapezoidal rule. Precision–recall curves are constructed from precision and recall values across thresholds, and the area under the precision–recall curve (AUPRC) is also computed using trapezoidal integration. AUPRC is particularly informative for evaluating models trained on imbalanced datasets, where the positive class is underrepresented.

### 2.6. Feature construction for the use case on predicting immunotherapy response from transcriptomics

#### GSVA pathway features

Pathway-level features were computed using Gene set variation analysis (GSVA) using the *GSVA* R package (v1.50.5). KEGG pathway gene sets were obtained from the *msigdbr* R package (v7.5.1), using the C2:CP:KEGG collection for Homo sapiens. Gene symbols associated with each pathway were grouped to construct gene sets. Gene set enrichment scores were then calculated for each sample using the GSVA algorithm with a Gaussian kernel cumulative distribution function, appropriate for normalized continuous expression data. GSVA transforms the gene expression matrix (genes × samples) into a sample-wise pathway activity matrix, where each value represents the relative enrichment of a pathway within a given sample.

#### K-medoids gene cluster features

Gene-level clustering features were derived using the k-medoids clustering algorithm implemented in the *cluster* R package (v2.1.4) via the CLARA (Clustering Large Applications) method. Genes were partitioned into k = 50 clusters based on their expression profiles across samples. For each cluster, a representative feature was computed by taking the mean expression value of all genes assigned to that cluster for each sample. This produced a reduced feature matrix where each column corresponds to the average expression of a gene cluster. During training, cluster assignments were estimated from the training data and subsequently reused when computing features for new datasets to ensure consistency between training and validation data.

#### Weighted Gene Correlation Network Analysis (WGCNA) module features

Co-expression module features were computed using the *WGCNA* R package (v1.73) (Langfelder P & Horvath S, 2008). The gene expression matrix was transposed to obtain a samples × genes format, and a weighted gene co-expression network was constructed using a soft-thresholding power of 6. Modules of co-expressed genes were identified using the blockwiseModules function with an unsigned topological overlap matrix (TOM) and a minimum module size of 30 genes. Each module was then summarized using its module eigengene, defined as the first principal component of the expression matrix of genes belonging to that module. This eigengene represents the dominant expression pattern of the module across samples. As with the clustering-based features, module assignments learned from the training data were reused to compute module eigengenes for external datasets.

## 3 Results

*pipeML* implements repeated and stratified k-fold cross-validation, optional model stacking with meta-learners, and cohort-aware validation strategies such as leave-one-dataset-out (LODO) analysis. Feature selection can be performed using repeated Boruta runs to identify robust predictors, while model training leverages the caret, tidymodels, parsnip and censored ecosystems packages for hyperparameter tuning and performance optimization across multiple classification and survival algorithms. *pipeML* further supports customized cross-validation fold construction functions, enabling feature engineering steps that must be repeated within each fold, as well as tunable feature-generation parameters that can be optimized jointly with model hyperparameters. Finally, the package provides built-in tools for prediction, threshold optimization, performance visualization (RO, PR and Kaplan-Meier curves), and model interpretation through variable importance via SHAP values, making it suitable for transparent and reproducible machine learning analyses in translational and systems biology studies.

### 3.1. Model selection and pipeline construction for classification and survival tasks

To demonstrate the application of *pipeML* for classification tasks, we applied it to the Breast Cancer Wisconsin Dataset (Wolberg et al. 1990; Zhang J. 1992; Blake CL, et al. 1998) available in the *mlbench* R package (v2.1.6). This dataset contains clinical measurements from breast tumor samples and is commonly used for evaluating binary classification algorithms. The predictors include nine cytological characteristics computed from digitized images of fine needle aspirates of breast masses, including clump thickness, uniformity of cell size and shape, marginal adhesion, epithelial cell size, bare nuclei, bland chromatin, normal nucleoli, and mitoses. The target variable represents tumor diagnosis, with two classes: benign and malignant.

Prior to model training, a binary target variable was created, where malignant tumors were encoded as 1 and benign tumors as 0. For example purposes, the dataset was subsequently divided into training and test sets using a stratified partition, with 70% of the observations used for model training and the remaining 30% reserved for testing.

Using *pipeML*, we performed automated model training and evaluation on the training dataset. Model selection was conducted using repeated k-fold cross-validation (5 folds with 10 repetitions) to obtain robust performance estimates and assess model stability (***Supplementary Figure 1***). Candidate models were evaluated using the AUROC as the primary performance metric, and hyperparameters were tuned automatically within the pipeline procedure. ***Supplementary Figure 2*** shows the RO and PR curves generated using the ML algorithm with the highest metric defined (in this case AUROC) (***Supplementary Table 1***).

For survival analysis tasks, we applied *pipeML* to the lung dataset (Loprinzi CL, et al. 1994) from the *survival* R package (v3.8.3). This dataset contains 228 patients with advanced lung cancer, including clinical variables such as age, sex, ECOG performance score, and other measurements relevant to prognosis. Prior to analysis, observations containing missing values were removed. The survival outcome is defined by the time to event (time) and the censoring indicator (status, 1 = death, 0 = censored). The dataset was subsequently divided into training and test sets using a stratified partition based on the event indicator, with 70% of the observations used for training and 30% reserved for testing.

Using *pipeML*, automated model training and evaluation were performed on the training data. Model selection was carried out using repeated k-fold cross-validation (5 folds with 10 repetitions) to obtain robust estimates of model performance and stability. Hyperparameters for candidate models were tuned automatically during training, and model performance was evaluated using the C-index. Similar to the classification tasks, ***Supplementary Figure 3*** shows the median C-index performance across folds for each survival algorithm ***(Supplementary Table 1***).

Both models, classification and survival were analyzed using SHAP values to retrieve the feature importance for the final model obtained from the *pipeML* training procedure, using all the cross-validation folds (***see Methods***).

### 3.2. *pipeML* achieves comparable median AUROC and AUPRC to standard machine learning frameworks

We compared the predictive performance of common machine learning models across three frameworks: *pipeML*, H2O AutoML (v3.44.0.3), and a Python scikit-learn implementation. Using the Sonar dataset (Blake CL, et al. 1998; Gorman RP, et al. 1998) from the R package *mlbench* (comprising 111 positive cases and 97 negative cases), we evaluated three models representing different families: GLM (Generalized Linear Model) as the linear model, Random Forest (RF) as a nonlinear ensemble tree-based method, and XGBoost as a gradient boosting algorithm capable of high predictive performance. Median AUROC and AUPRC were assessed over repeated cross-validation experiments (5 folds with 10 repetitions). *pipeML* achieved comparable median performance to both H2O AutoML and scikit-learn across all model families. Error bars, representing the median absolute deviation, indicate consistent performance across repeats (***Supplementary Figure 4***). These benchmarking results demonstrate that *pipeML* provides reliable predictions and in accordance with well-established machine learning pipelines.

### 3.3. *global dataset features* inflated predictive performance estimate under cross-validation training settings

To evaluate the impact of global dataset feature construction on model performance estimation, we designed a controlled experiment using the Sonar benchmark dataset. We generated a set of *global dataset features* by clustering original input features based on their pairwise correlations across samples and aggregating each cluster into a single feature via within-cluster averaging. This procedure yielded correlation-informed representations that depend on the global structure of the dataset.

We compared two cross-validation (CV) strategies. In the global feature construction CV setting, hereafter referred to as ‘standard CV’, *global dataset features* were computed once using the full dataset prior to model training and evaluation. In contrast, in the cross-validation fold-wise feature construction CV setting, hereafter referred to as ‘custom CV’, feature construction was embedded within the cross-validation loop using *pipeML*, such that correlation structure and feature aggregation were recomputed independently within each training cross-validation fold and then applied to the corresponding test samples, ensuring no leakage of information contributing to feature construction.

Across a diverse set of machine-learning models, standard CV consistently yielded higher median AUROC values compared to the custom CV strategy. Moreover, standard CV exhibited systematically lower variability across cross-validation folds, as measured by the median absolute deviation (MAD). In contrast, the custom CV approach resulted in reduced and more heterogeneous performance estimates, reflecting the removal of information leakage arising from *global dataset features* construction (***Supplementary Figure 5***). These results demonstrate that when new ‘aggregated’ features are derived from the original features using the global dataset structure, performing feature construction outside the cross-validation loop can lead to optimistically biased performance estimates. Properly including *global dataset features* construction within cross-validation is therefore important for reporting unbiased and reproducible estimates of predictive performance.

### 3.4. Use case: predicting immunotherapy response using public melanoma cohorts

To illustrate the utility of our pipeline and highlight the caveats of data leakage when computing *global dataset features* prior to model training in a real-world scenario, we applied our workflow to predict outcomes in six independent melanoma cohorts profiled by bulk RNA-sequencing, with gene expression matrices structured as genes × samples and annotated for immunotherapy response (***Supplementary Table 2***).

Using multiple independent cohorts allows us to reduce the risk that model performance is driven by characteristics specific to a single dataset, thereby providing a more reliable assessment of generalization. In particular, incorporating several cohorts helps mitigate cohort-specific technical and biological biases, while enabling evaluation of whether a no-data-leakage pipeline yields more realistic estimates of predictive performance across studies. To further ensure robustness, we adopted a Leave-One-Dataset-Out (LODO) validation strategy. In each iteration, models were trained on a subset of cohorts and evaluated on a held-out cohort that was not used during training. This design ensures that the test set remains fully independent and allows us to assess how well the trained models generalize to unseen cohorts, while minimizing biases arising from cohort-specific effects.

We started by filtering our gene expression matrix to retain the 5000 most variable genes across the full combined cohort. Within each LODO iteration, multiple feature representations were derived, including pathway activity scores using GSVA, gene cluster features obtained via k-medoids clustering, and co-expression module features derived from WGCNA (see Methods). We choose these approaches as they are commonly used in bioinformatics pipelines, and they tend to inadvertently include data-leakage during ML training. We compared the two cross-validation strategies (custom and standard CV) that differ based on when feature construction is performed relative to cross-validation splitting.

In the standard pipeline, features were computed once using the entire set of training cohorts prior to model training. The resulting feature matrices were then provided directly to *pipeML*, and cross-validation was applied to these precomputed features during model training and hyperparameter tuning. For the held-out cohort, features were computed independently using the same transformation procedure, or by projecting the data using parameters learned from the training cohorts (***see Methods; Supplementary Table 3***).

In contrast, the custom CV strategy integrates feature construction within the cross-validation procedure, implemented in *pipeML*. In this setting, gene expression data is provided directly to the training pipeline, and feature extraction is recomputed independently within each training fold during cross-validation. Specifically, GSVA enrichment scores, k-medoids gene clusters, and WGCNA co-expression modules are derived using only the samples belonging to the training portion of each fold. The learned feature transformations (e.g., cluster assignments or module definitions) are then applied to the corresponding validation samples within that fold. This design ensures that feature construction does not use information from validation samples during model training, thereby preventing potential data leakage (***Supplementary Table 4***).

Predictive performance was evaluated on the held-out LODO cohort using AUROC, enabling comparison between the standard and custom CV strategies in terms of their ability to generalize to independent datasets ***(Supplementary Table 5 and 6***).

Models trained on the WGCNA features using the custom CV strategy showed significantly lower AUROC values relative to the standard CV across all cohorts (***Figure 2, top***). This difference reflects the removal of information leakage that occurs when *global dataset features* are computed using the full dataset prior to CV. Model prediction, on the contrary, showed consistent performance estimates with the AUROC observed on held-out cohorts during training for the custom CV strategy, whereas standard CV tended to significantly reduce its performance compared to what was shown during training. Notably, the discrepancy was most evident for the Hugo cohort, where standard CV suggested higher predictive performance during training than what was observed during external validation (***Figure 2, bottom***). Similar patterns were observed for GSVA enrichment scores and k-medoids gene clusters. In these cases, standard CV generally produced higher training AUROC values than the custom strategy, but performance during testing was either comparable between the two approaches or lower for the standard CV in some cohorts (***Supplementary Figure 6***).

**Figure 2.**
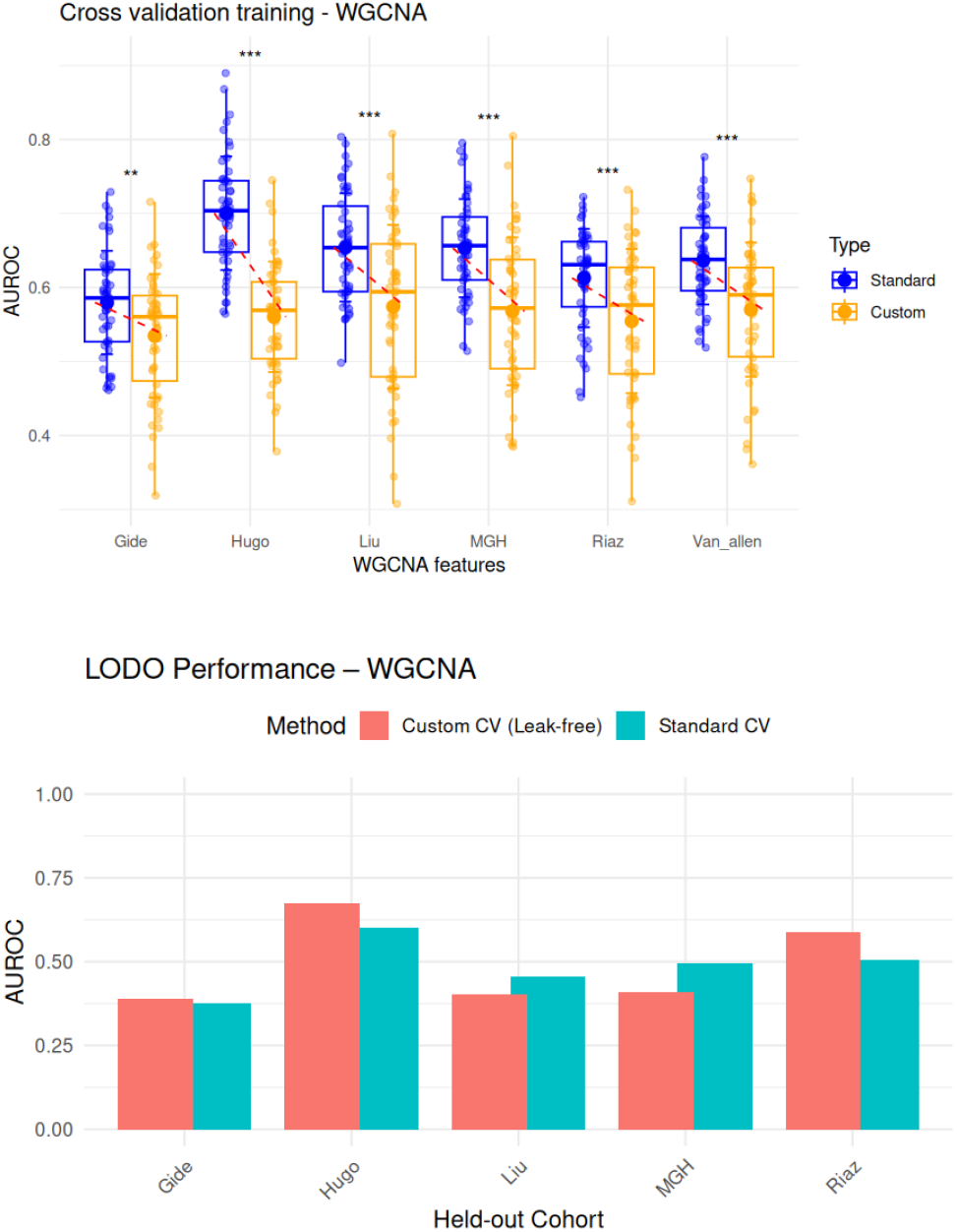
AUROC predictive performance for held-out cohorts comparing standard cross-validation (CV) and custom (leak-free) CV strategies using WGCNA features. (top) Boxplots of AUROC per cohort for each CV strategy across all models; paired differences were assessed using a paired t-test. Statistical significance is indicated as follows: *** p < 0.001, ** p < 0.01, * p < 0.05, ns = not significant. (bottom) Median AUROC for each cohort using standard and custom CV strategies, highlighting overall performance differences between CV methods.

Overall, these results indicate that recomputing global features within each CV fold reduces information leakage and yields performance estimates that better reflect model generalization to independent datasets.

### 3.5. Prediction performance is highly affected by the choice of hyperparameters during construction of custom CV folds

Prediction performance can be strongly influenced by the choice of hyperparameters used during the construction of custom CV folds. Custom fold construction functions may involve several user-defined parameters, whose optimal configuration depends on the dataset, the analytical objective, and the predictive metric of interest. To address this, *pipeML* provides a framework that enables systematic tuning of these hyperparameters and optimization of the fold construction procedure based on performance metrics such as AUROC, AUPRC, or the concordance index (C-index).

Similar to traditional hyperparameter tuning in machine learning models, *pipeML* evaluates multiple parameter combinations, calculates predictive performance across CV resamples, and identifies the configuration that yields the best performance. In this framework, both the parameters governing the custom fold construction and the parameters of the machine learning model can be optimized jointly.

To illustrate this functionality, we applied the WGCNA algorithm (see Methods), which includes several tunable parameters such as the soft-thresholding power, minimum module size, module merging threshold, and module splitting sensitivity. For simplicity and computational efficiency, this analysis was restricted to the Hugo cohort (Hugo et al. 2016), using the 5,000 most variable genes as input features. Specifically, we evaluated combinations of soft-thresholding power (6, 8, 10), minimum module size (20, 50, 100), module merging threshold (0.15, 0.25, 0.35), and module splitting sensitivity (1, 2, 3), resulting in 81 possible WGCNA parameter configurations.

For each configuration, predictive models were trained using logistic regression with elastic-net regularization, where the mixing parameter α (values 0 and 1) and the regularization parameter λ (20 values between 0.001 and 1) were also tuned. The joint exploration of WGCNA and machine learning hyperparameters resulted in a total of 3,240 evaluated parameter combinations (***Supplementary Table 7***).

During cross-validation, *pipeML* computes features derived from each WGCNA configuration, trains predictive models across the machine learning parameter grid, and evaluates model performance across resamples. This process allows the identification of the parameter combination that maximizes the selected performance metric.

***Supplementary Figure 7*** summarizes the resulting predictive performance (AUROC) across all tested parameter combinations. Each point represents the median AUROC achieved by a specific combination of WGCNA and logistic regression hyperparameters, illustrating how variations in both feature construction and model regularization influence predictive performance. The highest AUROC (0.83) was achieved using an elastic-net mixing parameter α = 1 and a regularization parameter λ = 0.106, combined with WGCNA parameters power = 8, minimum module size = 20, module merging threshold = 0.15, and module splitting sensitivity (deepSplit) = 1. This configuration (***highlighted in red***) was automatically selected by *pipeML* to train the final predictive model.

## 4 Discussion

*pipeML* provides a flexible and robust framework for machine learning analyses in biological settings where features are dataset-dependent and traditional cross-validation strategies can introduce information leakage. By enabling custom cross-validation fold construction and fold-aware feature recomputation, *pipeML* ensures that model training and evaluation reflect realistic predictions on unseen data.

The importance of realistic estimations of performance in the cross-validation setting cannot be overstated: this range is an estimate of the best- or worst-case scenario performance and, since we never know on which external dataset the model could be run, these estimates correspond to the bounds that we could expect on the performance in independent datasets, assuming there are no major differences between the original and external datasets. If the cross-validation performance estimate is generally too high, we might drastically overestimate the low-bound for the performance of the predictor in practical applications.

A main limitation of this approach is that this realistic estimation of expectations on the model performance comes at a cost: often the calculation of *global dataset features* rely on approaches such as statistical aggregation or correlation, that normally require a large number of samples to generate robust features. Performing feature construction separately in each fold will necessarily reduce the number of samples used and, hence, worsen the robustness of the calculated features, especially when sample numbers are low. These aspects should be considered when choosing the cross-validation strategy, based on the exact goal for making the model in the first place.

*pipeML* allows users to integrate complex, biologically informed feature engineering steps, including for example enrichment-based or correlation-based features, without compromising model validity.

Implemented in R, *pipeML* seamlessly integrates with the Bioconductor ecosystem and existing bioinformatics workflows, supporting both classification and survival analyses. As predictive modeling in biomedical research increasingly relies on global dataset feature representations, *pipeML* offers a scalable, modular, and leakage-aware solution that delivers reliable performance estimates, while remaining fully adaptable to user-defined feature construction strategies and current machine learning workflows.

## Author contributions

M.H. conceived the project. M.H. developed the package and methods. M.H performed the benchmark and the analyses. V.P. and M.H wrote and reviewed the manuscript. V.P. supervised the project.

## Acknowledgements

The authors thank Raphaël Mourad and Hafida Hamdache for their helpful feedback on the package methods. We also thank Andrei Zinovyev for his valuable critical reading of the previous version of the manuscript and providing insightful comments.

## Data availability

No new experimental data was generated as part of this study. All the datasets used are publicly available as described in ***Supplementary Table 2***. Code to reproduce the figures of this paper can be found at https://github.com/VeraPancaldiLab/pipeML_paper

