## Supplementary Material for "A new pipeline for cross-validation fold-aware machine learning prediction of clinical outcomes addresses hidden data-leakage in omics based ‘predictors’"

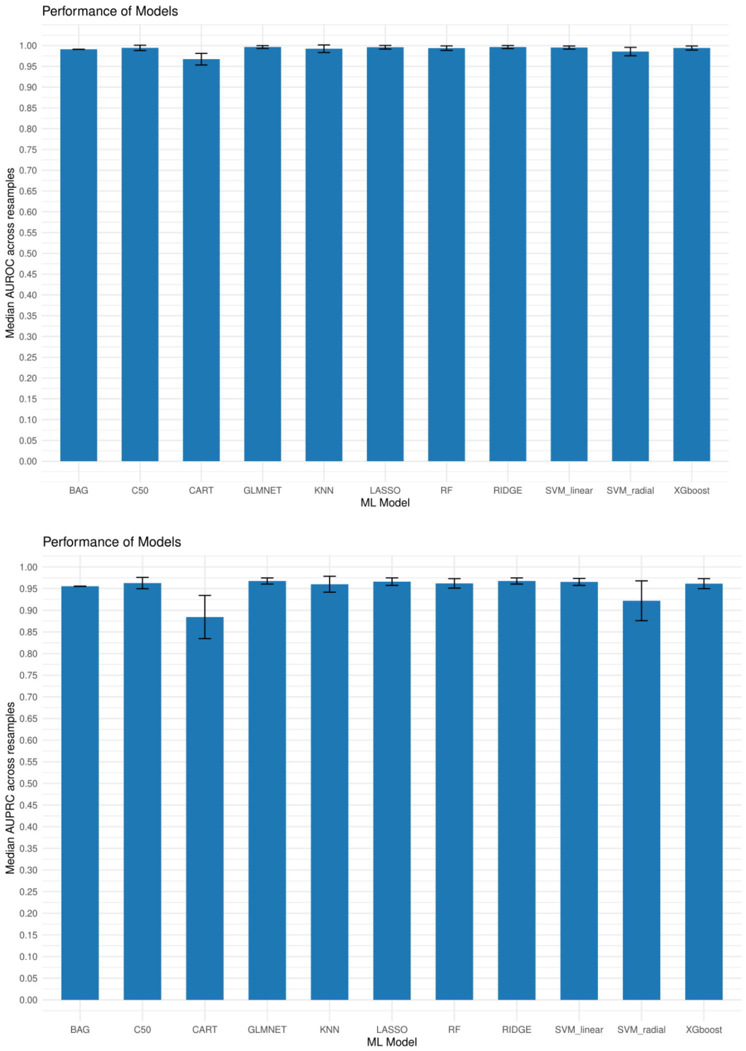


***Supplementary Figure 1.*** *Performance and interpretation of classification models trained on the Breast Cancer Wisconsin dataset using a 70% training and 30% test split.* ***(A)*** *Cross-validated AUROC values for different classification algorithms evaluated on the training set.* ***(B)***

*Cross-validated AUPRC values for different classification algorithms evaluated on the training set.*

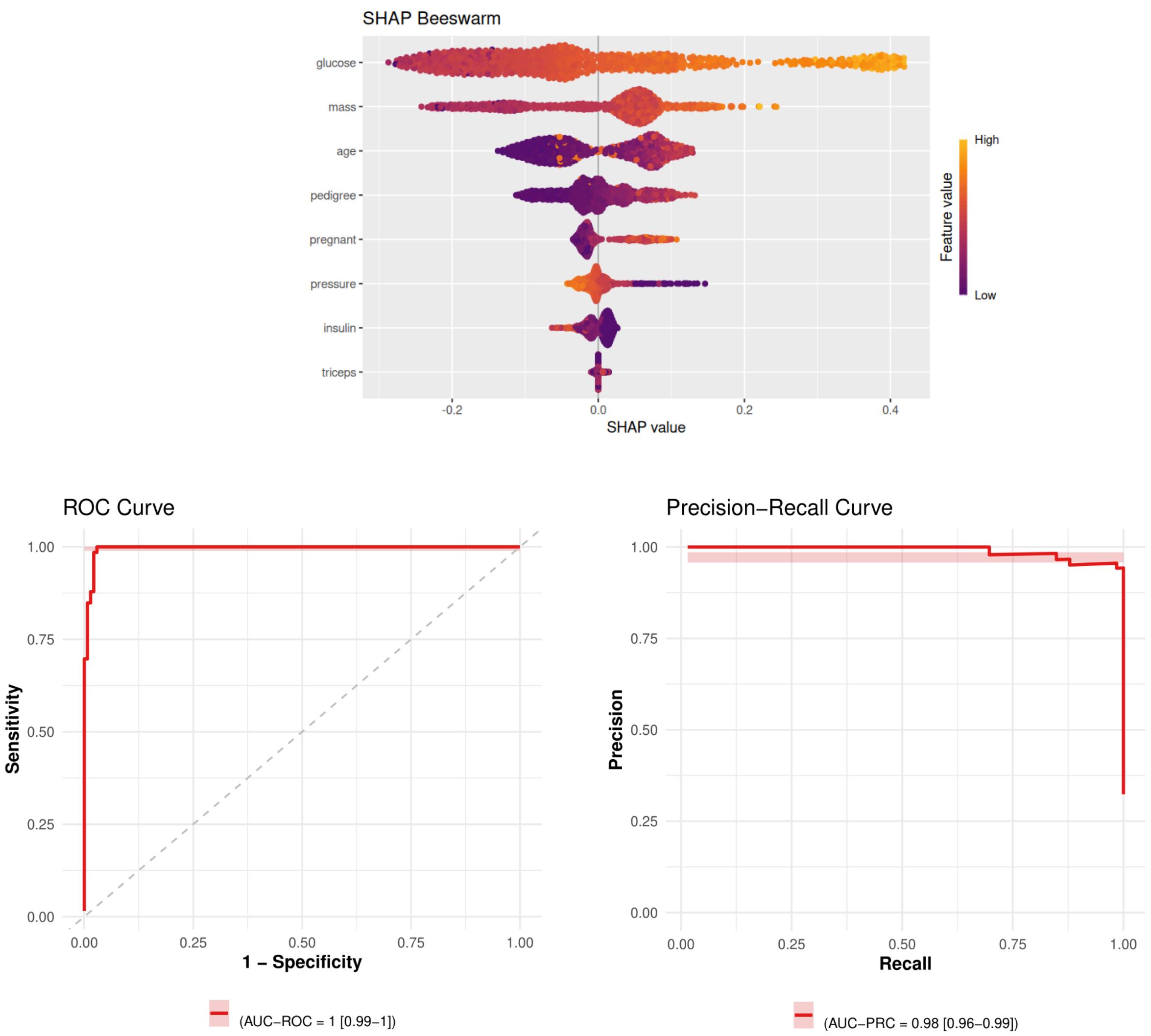


***Supplementary Figure 2.*** *Performance and interpretation of classification models trained on the Breast Cancer Wisconsin dataset using a 70% training and 30% test split.* ***(A)*** *SHAP values showing the relative contribution of predictive variables to the model predictions.* ***(B)*** *Receiver operating characteristic (ROC) and* ***(C)*** *precision–recall (PR) curves evaluating model performance on the independent test set.*

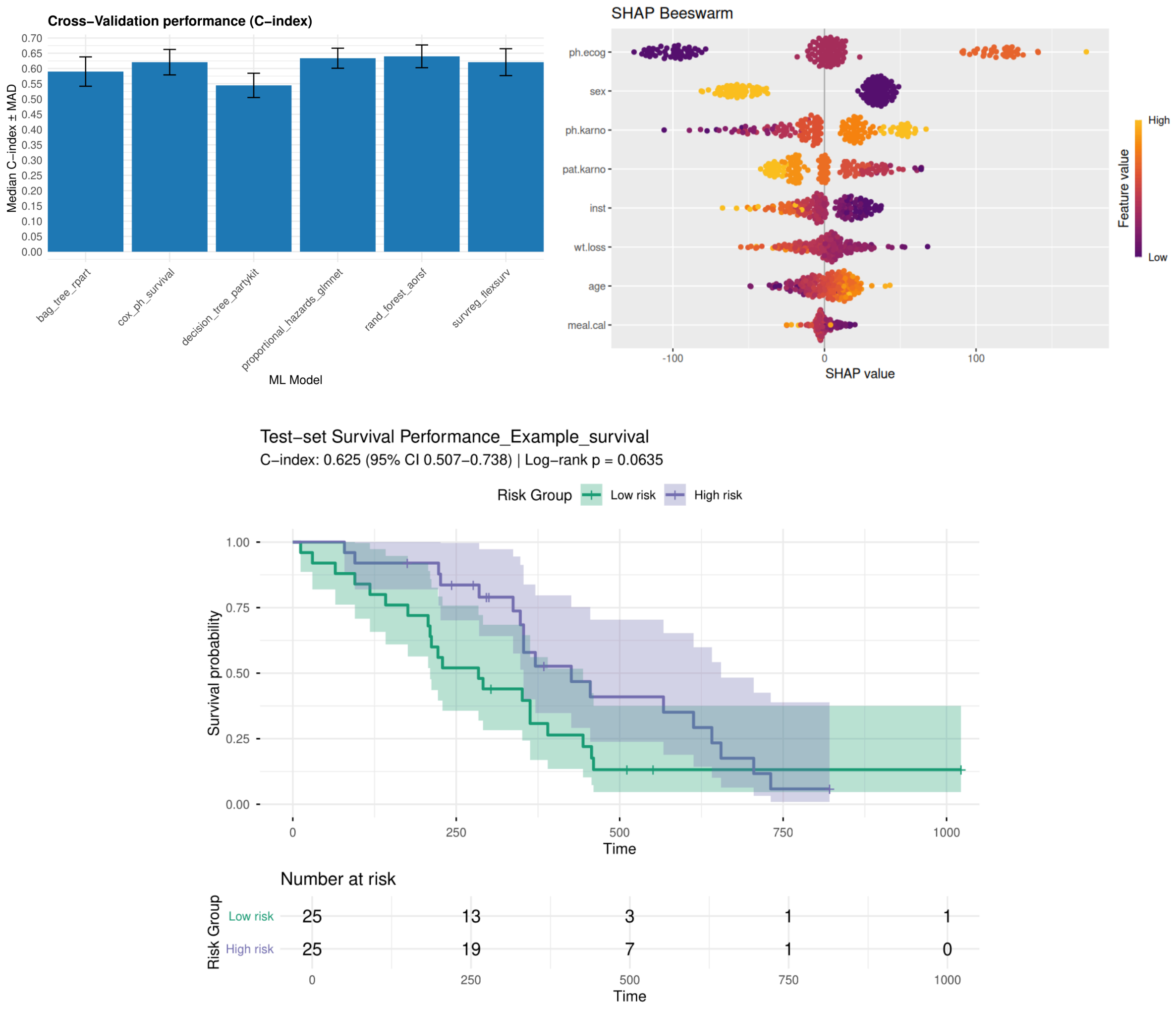


***Supplementary Figure 3.*** *Performance and interpretation of survival models trained using a 70% training and 30% test split.* ***(A)*** *Cross-validated concordance index (C-index) for different survival analysis algorithms evaluated on the training set.* ***(B)*** *SHAP values showing the contribution of predictive variables to the model predictions.* ***(C)*** *Kaplan–Meier survival curves illustrating risk stratification performance on the independent test set.*

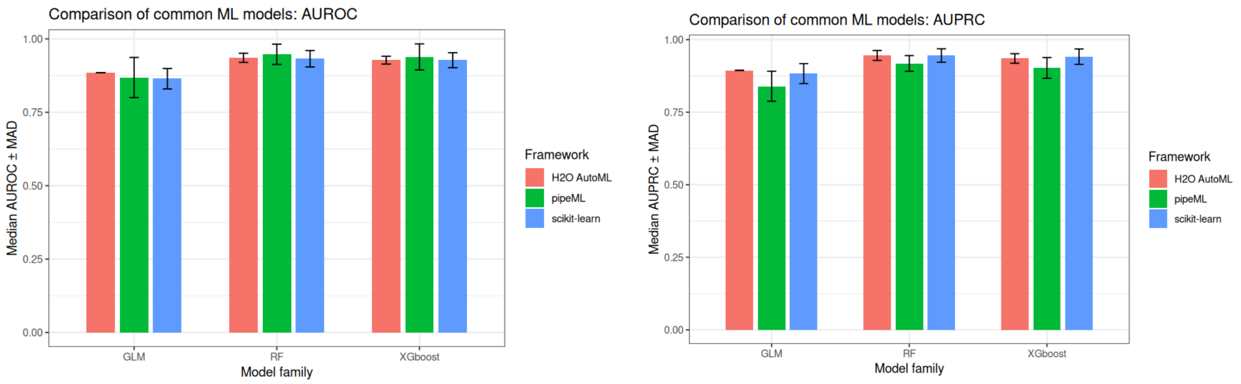


***Supplementary Figure 4.*** *Comparison of predictive performance between established machine learning frameworks and pipeML across three model families: generalized linear models (GLM), random forests (RF), and XGBoost.* ***(A)*** *Area under the receiver operating characteristic curve (AUROC).* ***(B)*** *Area under the precision–recall curve (AUPRC).*

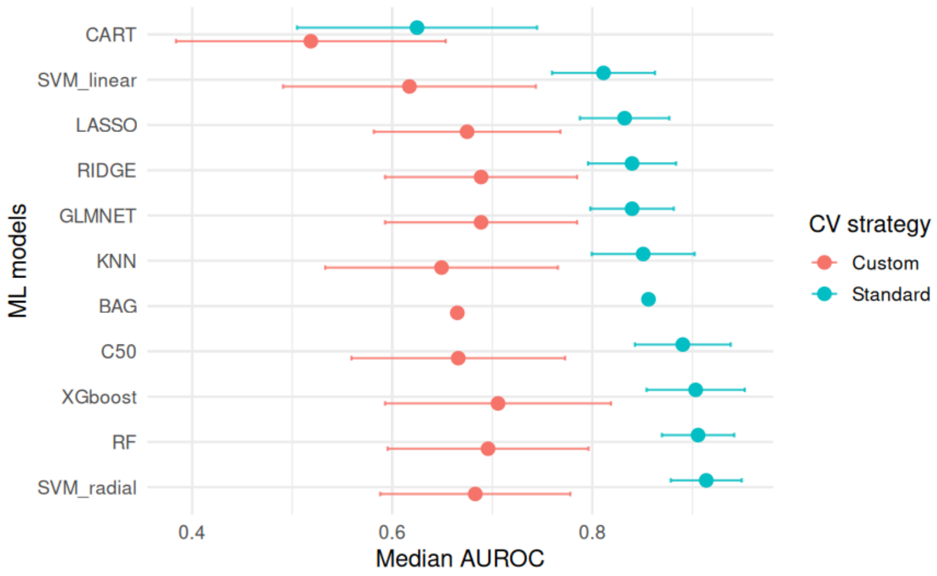


***Supplementary Figure 5.*** *Median AUROC comparing standard and custom cross-validation (CV) strategies across 11 machine learning models implemented in pipeML. Error bars indicate the median absolute deviation (MAD) across CV resamples.*

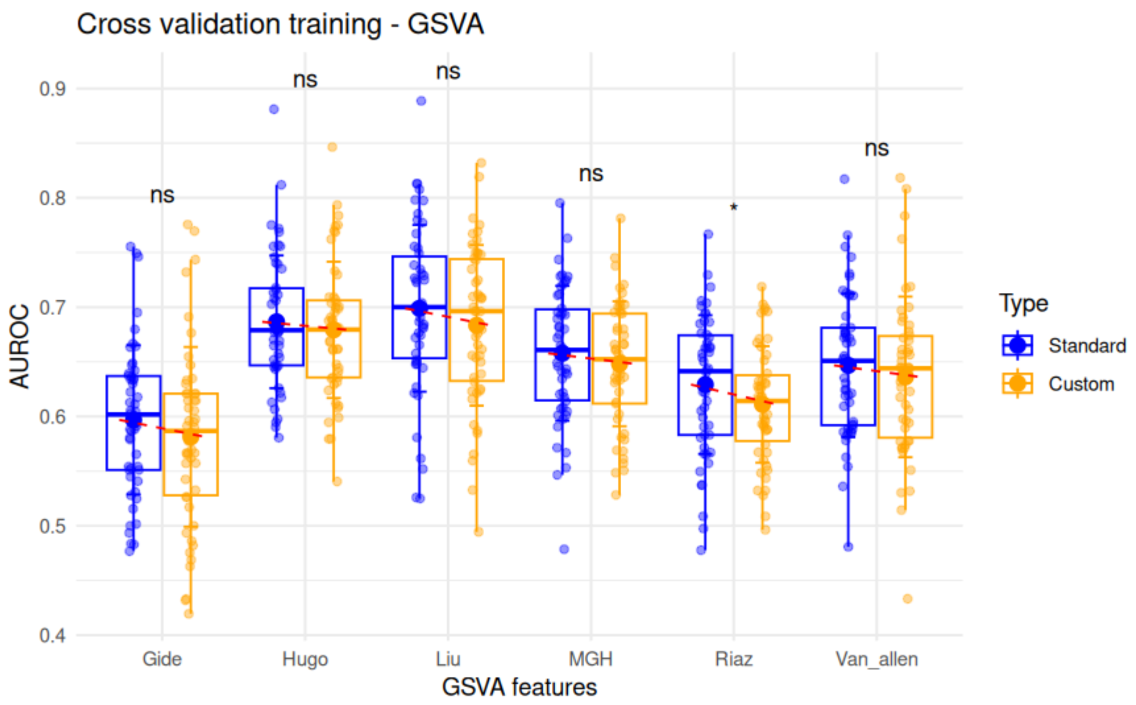
Cross validation training - GSVA
AUROC
0.9
0.8
0.7
0.6
0.5
0.4
ns
ns
ns
ns
*
ns
Gide
Hugo
Liu
MGH
Riaz
Van_allen
GSVA features
Type
Standard
Custom

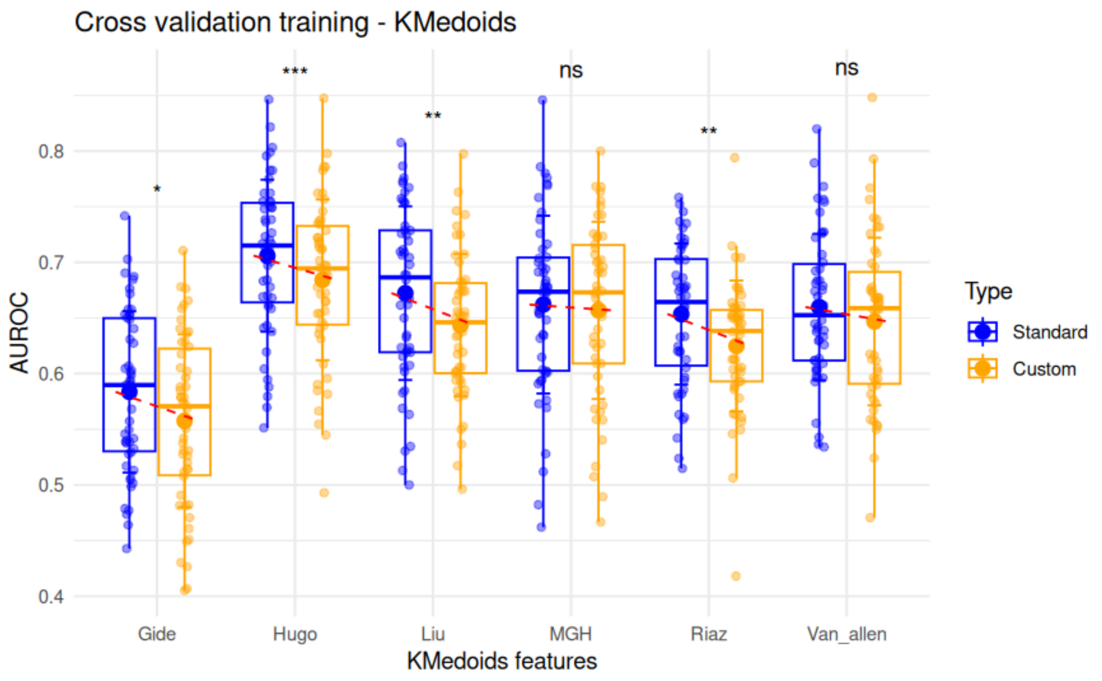
Cross validation training - KMedoids
AUROC
0.8
0.7
0.6
0.5
0.4
*
***
**
ns
**
ns
Gide
Hugo
Liu
MGH
Riaz
Van_allen
KMedoids features
Type
Standard
Custom

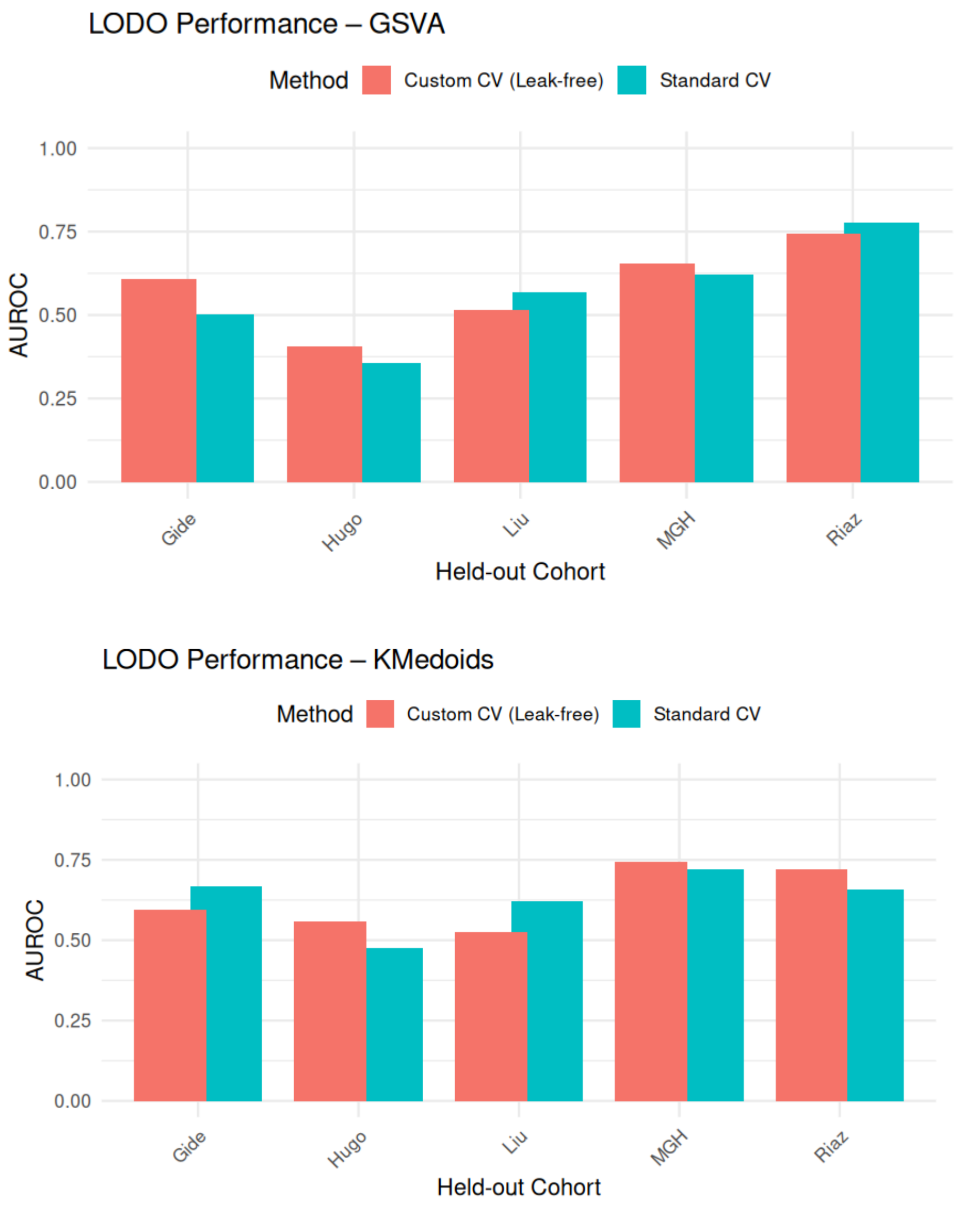


***Supplementary Figure 6.*** *AUROC predictive performance for held-out cohorts comparing standard cross-validation (CV) and custom (leak-free) CV strategies.* ***(A–B)*** *Boxplots of AUROC per cohort for each CV strategy across all models using GSVA* ***(A)*** *and KMedoids* ***(B)*** *features. Paired differences were assessed using a paired t-test. Statistical significance is indicated as follows: *** $p < 0.001$, ** $p < 0.01$, * $p < 0.05$, ns = not significant.* ***(C–D)*** *Median AUROC for each cohort using standard and custom CV strategies for GSVA* ***(C)*** *and KMedoids* ***(D)*** *features, highlighting overall performance differences between CV methods.*

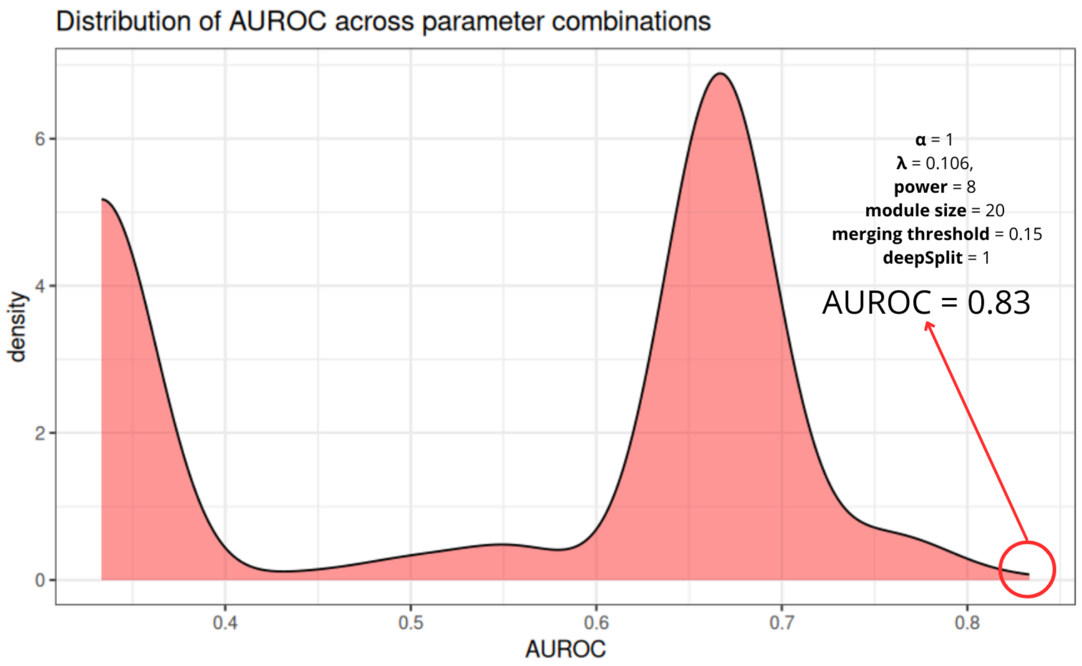


***Supplementary Figure 7.*** *Model performance (AUROC) across 3,240 hyperparameter combinations evaluated using custom cross-validation (CV) folds. Each point represents the median AUROC obtained for a specific hyperparameter configuration. The distribution of AUROC values across all configurations is shown as a density plot, illustrating the overall variability in predictive performance resulting from different hyperparameter settings. Highlighted in red is the parameter combination which maximizes the AUROC across training.*

**Supplementary Tables are provided as .xlsx files**

**Supplementary Table 1:** Training and prediction metrics obtained for example datasets (Result 1) including AUROC, AUPRC, hyperparameters, Sensitivity, Specificity, fpr, Accuracy, Precision, Recall, F1, MCC scores.

**Supplementary Table 2:** Melanoma cohorts information used in use-case results.

**Supplementary Table 3:** Training metrics obtained for use case example per held-out cohort and feature set (GSVA, KMedoids and WGCNA) including AUROC, AUPRC, hyperparameters, Sensitivity, Specificity, fpr, Accuracy, Precision, Recall, F1, MCC scores for *standard* k-fold strategy.

**Supplementary Table 4:** Training metrics obtained for use case example per held-out cohort and feature set (GSVA, KMedoids and WGCNA) including AUROC, AUPRC, hyperparameters, Sensitivity, Specificity, fpr, Accuracy, Precision, Recall, F1, MCC scores for *custom* k-fold strategy.

**Supplementary Table 5:** Prediction metrics obtained for use case example per held-out cohort and feature set (GSVA, KMedoids and WGCNA) including AUROC, AUPRC, hyperparameters, Sensitivity, Specificity, fpr, Accuracy, Precision, Recall, F1, MCC scores for *standard* k-fold strategy.

**Supplementary Table 6:** Prediction metrics obtained for use case example per held-out cohort and feature set (GSVA, KMedoids and WGCNA) including AUROC, AUPRC, hyperparameters, Sensitivity, Specificity, fpr, Accuracy, Precision, Recall, F1, MCC scores for *custom* k-fold strategy.

**Supplementary Table 7:** AUROC and AUPRC performance across the different hyperparameters combinations using WGCNA as custom fold construction function.
